# Jasmonates regulate auxin-mediated root growth inhibition in response to rhizospheric pH in Arabidopsis

**DOI:** 10.1101/2023.08.22.554241

**Authors:** Ajit Pal Singh, Rajni Kanwar, Ajay K. Pandey

**Author notes:** Authors for correspondence Ajit Pal Singh; Ajay K Pandey.

## Abstract

Rhizospheric pH severely impacts plant growth and fitness through a numerous process and has emerged as a major determinant of crop productivity. Despite numerous attempts, the key questions related to plants response against rhizospheric pH remains largely elusive. The present study provides a mechanistic framework for rhizospheric pH-mediated root growth inhibition (RGI). Utilizing various genetic resources combined with pharmacological agents and high-resolution confocal microscopy, the study provides direct evidences for the involvement of jasmonates and auxin in rhizospheric pH-mediated RGI. We show that auxin maxima at root tip is tightly regulated by the rhizospheric pH. In contrast, jasmonates (JAs) abundance inversely correlates with rhizospheric pH. Further, JA mediated regulation of auxin maxima by *GRETCHEN HAGEN 3* (*GH3*) family genes explains the pattern of RGI observed over the entire range of rhizospheric pH. Our findings revealed auxin as the key regulator of RGI during severe pH conditions, while JAs antagonistically regulate auxin response against rhizospheric pH.

**Highlight:** The current study identifies the mechanistic framework of rhizospheric pH mediated root growth inhibition in model plant Arabidopsis through a prominent crosstalk between two phytohormones i.e. auxin and jasmonates.

## Introduction

Rhizospheric pH (pH_Ext_) is one of the most important environmental factors affecting plant growth and physiology (**Fageria and Zimmermann, 1998**). Most plants only survive over a very narrow range of pH_Ext_ by maintaining a relatively constant cytoplasmic pH ranging from 5.8-6.3 (**Arnon and Johnson, 1942**). It has been observed that soil pH is being disturbed at an alarming rate resulting into unfit soil for cultivation. For an estimate, around 40% of the world’s arable land has acidic soil and around 1 billion ha is affected by salts, with around 60% of this having alkaline pH (**Zhang et al., 2023**). Change in pH_Ext_ not only impact the rhizospheric microbiome but also have a profound effect on nutrient availability, root exudates and plant physiology. Lower pH_Ext_ below pH_5.5_ (acidic soils) results in heavy-metal toxicity, decreased nodulation and nitrogen fixation etc., while pH_Ext_ higher than pH_6.5_ (alkaline soils) results in decreased Fe, Mn, Zn, B and phosphate bio-availability for plants (**Tsai & Schmidt, 2021**). Being sessile, plants need to compromise their defense strategies, nutrient homeostasis, metabolite fluxes, and transcriptional profiles to adapt against the changing rhizospheric pH (**Lager et al., 2010; Tsai and Schmidt, 2021; Bailey et al., 2022**).

Though plants are known to respond to pH_Ext_ by reduced yield, their ability to perceive pH_Ext_ is a heavily debated question. Recent findings suggest that plants not only perceive pH_Ext_ and also respond to it possibly through trans-acting factors (**Tsai and Schmidt, 2021**). Generally, low pH_Ext_ is proposed to favour plant growth by cellular expansion (**Cosgrove, 1999**). This proposed hypothesis never fit into the actual scenario where both acidic as well as alkaline pH_Ext_ are reported to result in root growth inhibition (**Duan et al., 2023; Jain and Schmidt, 2023**). However, the dependency of cell elongation on pH indicates a perfect relation between root growth and pH_Ext_ (**Barbez et al., 2017**). Though auxin-regulated root growth is supported by acid growth theory; it remained the subject of debate for its dual nature of being stimulatory as well as inhibitory (**Moloney et al., 1981; Lado et al., 1976; Sauer et al., 2013; Fendrych et al., 2018**). Based on the transcriptional response, recent studies have hinted at the involvement of auxin signalling machinery in low pH_Ext_ mediated root growth inhibition (**Lager et al., 2010, Bailey et al., 2022**). Therefore, an extensive study of root growth inhibition during varying pH_Ext_ conditions is required to understand the mechanistic aspect of soil pH perception and response.

The present study explores the role of phytohormones for pH_Ext_ mediated root growth inhibition, where we show that auxin maxima at root tip co-linearly associated with the entire range of pH_Ext_. While auxin seems to regulate root growth inhibition (RGI) by its low and high abundance at the root tip during acidic and alkaline pH conditions, respectively, it may not have the pH sensing role. Though, a recent report (**Duan et al., 2023**) suggested the role of polar auxin transport in the alkaline pH mediated root growth inhibition. However, it is unclear if pH_Ext_ mediated RGI is regulated at the level of auxin polar transport, its biosynthesis or catabolism. Our work here, provides definite evidences to support biosynthesis and catabolism mediated auxin accumulation during varied pH_Ext_ range. Further, we show that JA (jasmonate acid) abundance/signalling is inversely associated with the pH_Ext_ value and works upstream of auxin by negatively regulating auxin abundance at the root tip. Therefore, this study while considering a range of rhizospheric pH provides a conceptual framework for root growth inhibition in response to rhizosphere pH.

## Materials and Methods

### Plant material and growth conditions

The seed stocks used in this study are Arabidopsis WT (ecotype Col-0 or Col-gl, as indicated in the text), *coi1-16* (CS67817), *jar1-11* (CS67935), *tir1-1* (CS3798), DII:VENUS (CS799173), DR5rev::GUS (**Friml et al., 2003**), p35S::Jas9-N7-VENUS (**Larrieu et al., 2015**), pCYCB1;1::GUS (**Ferreira et al., 1994**), pPIN1::PIN1-GFP, pPIN2::PIN2:GFP and pPIN7::PIN7:GFP (**Blilou et al., 2005; Friml et al. 2003).**

Seeds were surface sterilized with 2% sodium hypochlorite for 5 min followed by 5 times wash (each 5 min) with autoclaved water. After surface sterilisation seeds were stratified at 4℃ for 3 days and germinated on ½MS media (pH_5.8_) solidified with 0.8% agar. After 4 days of germination, seedlings were transferred to the indicated stress conditions. All the growth assays were performed at 22℃ with 16 h photoperiod.

### Root gravitropic response

Col-0 seeds were plated on ½MS media (square plates) with different pH conditions and allowed to germinate. After 4 days of normal growth on the same media plates were rotated by 90-degree angle (clockwise) so that the gravity vector is shifted by 90 degree. Images were captured after 10 hours of the re-orientation and root angle was measured using ImageJ along the initial gravity vector.

### Split root assay

For local and systemic sensing of pH_Ext_, equally germinated Col-0 seedlings (4 days old) were transferred to ½ MS-agar media with pH_5.8_ followed by cutting the root tip of each seedling 1 day after the transfer. After 6 days of root tip cut seedlings with appropriately bifurcated roots were selected and transferred to the combination of pH_Ext_ conditions as indicated. The starting point was marked on the same day of seedling transfer and root growth after 6 days of the treatment was considered for the relative root growth analysis.

### Treatments

Uniformly germinated 4 days old seedlings were transferred to ½ MS-agar media having different pH conditions either with or without MeJA (Methyl jasmonate), NAA (1-naphthaleneacetic acid), TIBA (2,3,5-triiodobenzoic acid), NPA (N-1-naphthylphthalamic acid), Kyn (L-kynurenine), PPBo (4-phenoxyphenylboronic acid) and BBo (4-biphenylboronic acid) with mentioned concentrations. Images were captured after 5 days of treatment. Root growth after onset of the treatment was considered for analysis.

Stocks used for the study are as follows: MeJA (10mM in DMSO), NAA (10mM in ethanol), TIBA (5mM in ethanol), NPA (2mM in DMSO), kyn (2mM in DMSO), BBo acid (4mM in DMSO), PPBo (4mM in DMSO). All these chemicals were purchased from sigma.

### Confocal microscopy

Uniformly germinated Arabidopsis seedlings (as mentioned) were transferred to the treatment as indicated and signals were observed in the root tip after 3 days of the treatment with an inverted confocal microscope (ZEISS LSM 880). Excitation and emission conditions for the dyes/fluorophores were set as followed: DR5rev::GFP; PIN1:GFP, PIN2:GFP, PIN7:GFP (Ex. 488nm, Em. 493-598nm), DII:VENUS (Ex. 514nm, Em. 517-605nm), Jas9:VENUS (Ex. 514nm, Em. 517-636nm) and propidium iodide (Ex. 561nm, Em. 566-718nm).

Propidium iodide staining was performed for visualising cell size by incubating seedlings in 15µM PI for 10 min in dark followed by 3 times was with distilled water. The onset of elongation was defined as the point where cortical cell length becomes twice of its preceding cell.

### GUS activity assay

CycB1;1:GUS signals were analysed in primary roots after 3 days of the indicated treatment in primary roots. Briefly, after treatment the seedlings were transferred to GUS buffer (containing 1mM 5-bromo-4-chloro-3-indolyl-β-glucuronide sodium salt in 50mM sodium phosphate buffer, pH 7.0, 10mM Na-EDTA, 0.5mM ferricyanide, 0.5mM ferrocyanide and 0.1% Triton X-100). Seedlings were incubated at 37 °C for overnight in dark conditions and images were captured using stereo-zoom microscope.

### Promoter analysis

Promoter (2Kb upstream of transcription start site) was identified for each gene and analysed using PlanPan2.0 (http://plantpan2.itps.ncku.edu.tw/) for all the cis elements present in the promoter. Only auxin and JA responsive cis elements were considered for the analysis after manual removal of duplicate entries.

### Gene expression analysis

Root tissue of the seedlings were collected after 8 days of the indicated treatment. Total RNA was extracted using TRIzol™ Reagent (Invitrogen™) followed by DNaseI digestion to get rid of gDNA. This RNA was then used for cDNA synthesis using High Capacity cDNA Reverse transcription Kit (Applied biosystems) following manufacturer’s instruction. For qRT-PCR, TB Green^TM^ Premix Ex Taq^TM^ (Takara) was used as per the manufacturer’s instructions. Fold change analysis was done using 2^-ΔΔCT^ method with *Actin2* (AT3G18780) as internal control gene for normalization. Primers were designed manually and are listed in **Table S1**.

### Statistics and data analysis

Total number of seedlings per line/conditions were considered as the sample size (n). For pair-wise statistical analysis, two-tailed student’s *t-*test was used with p-value < 0.05 using Microsoft Excel software. Line and bar graphs were generated using Microsoft Excel software and box-whisker plots were generated using R-4.3.1 program (https://cran.r-project.org/bin/windows/base/).

PRL (primary root length) was analysed using ImageJ (Fiji, https://imagej.net/software/fiji/downloads) considering appropriate scales. For normalisation, each PRL/fluorescence value was divided by the average PRL value at the control/optimal condition (as indicated along with each experiment). The average value of normalized PRL or fluorescence was used for plotting with SE among the replicates. All the experiments were performed at least thrice.

## Results

### Rhizospheric pH regulate primary root growth reversibly

The ability of plants to perceive rhizospheric pH (pH_Ext_) is a long-debated question, therefore, we aimed to clarify whether pH_Ext_ is perceived by plants and if so how do plants respond against different pH_Ext_ conditions. To have a detailed insight into the plant’s response against pH_Ext_, we analysed the impact of a range of pH_Ext_ on Arabidopsis root growth. Our observations marked rhizospheric pH_5.8_ supporting the best proliferating primary root while both the pH_Ext_ extremes (towards acidic and alkaline) inhibited the primary root growth. The degree of primary root length (PRL) inhibition correlated well with the degree of pH_Ext_ change from pH_5.8_. Therefore, in order to convey our findings, we selected pH_5.8_ as optimum pH_Ext_ and pH_3.8_ and pH_7.8_ as treatments for our entire study (**Figure 1a, Figure S1**). Interestingly, the RGI (root growth inhibition) caused by pH_3.8_ and pH_7.8_ was entirely reversible as it shows the comparatively similar growth rate when recovered with pH_5.8_ (**Figure 1b**).

**Figure 1.**
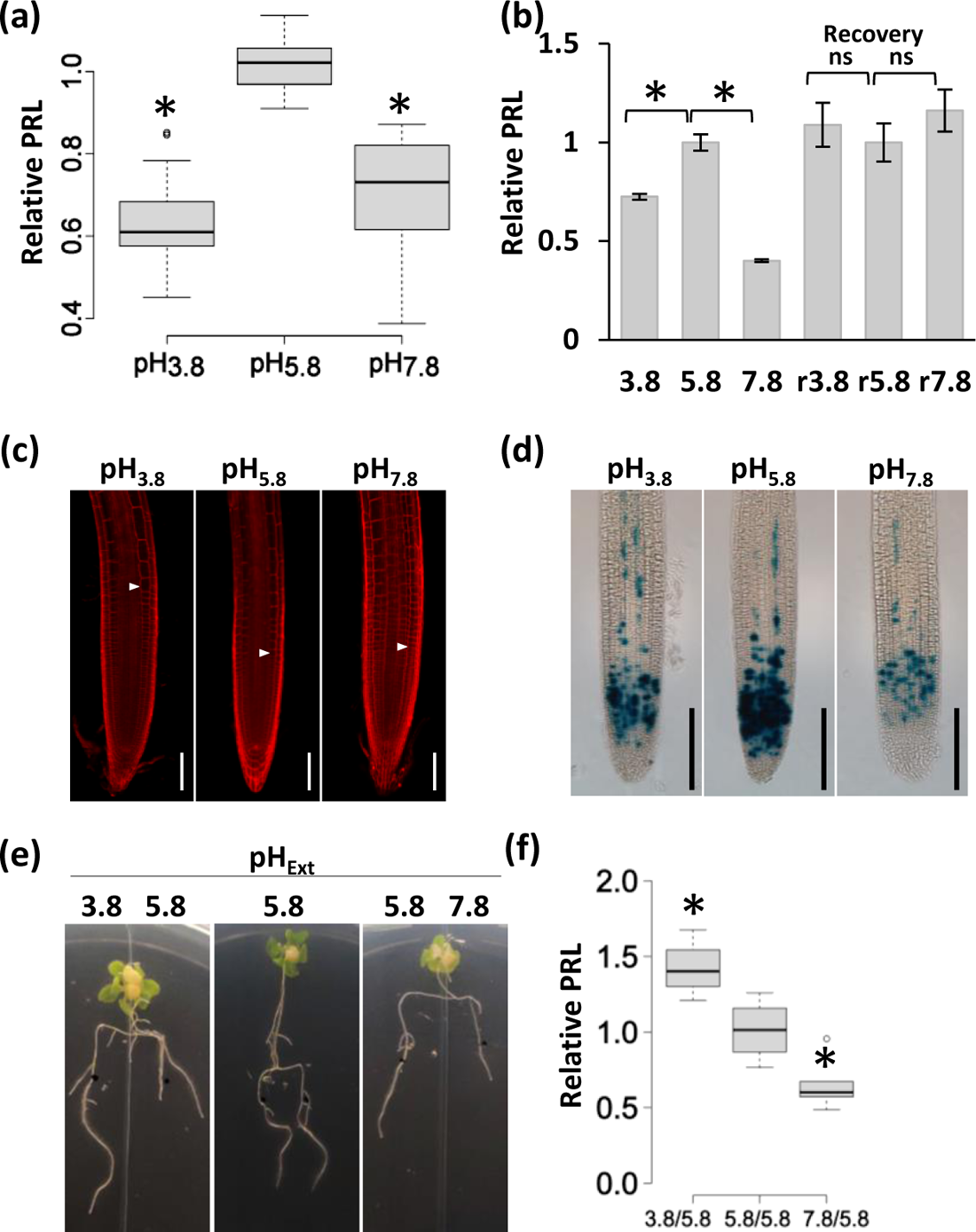
Rhizospheric pH regulate root growth inhibition in Arabidopsis. **(a)** The effect of rhizospheric pH on primary root length (PRL) in Col-0. PRL was analysed after 5 days of treatment and normalised with average PRL at pH_5.8_ for plotting. Y-axis: relative PRL on different pH_Ext_ conditions as compared to optimal rhizospheric pH_5.8_ (i.e. root length pH_Ext_/root length pH_5.8_), x-axis: pH_Ext_ values, n≥20. **(b)** Bar graph showing reversible nature of RGI in response to pH_Ext_. PRL was plotted relative to pH_5.8_ after pH_Ext_ treatment of 5 days. All the seedlings (irrespective of the treatment) were then recovered with pH_5.8_ (r5.8) for next 5 days and normalized values were plotted (r3.8 and r7.8) against pH_5.8_ (r5.8). Asterisks marks the significant difference compared with root length on pH_5.8_, student’s *t-* test, p-value < 0.05, n ≥ 10. **(c)** Representative image showing effect of pH_Ext_ on root cell files. Roots from seedlings grown on indicated pH_Ext_ for 3 days were treated with PI and visualized under confocal microscope. Cell length was analysed for first elongated cortical cell (marked with arrow). Scale bar=100µm. **(d)** Effect of pH_Ext_ on RAM activity. CYCB1;1:GFP seedlings were subjected to different pH_Ext_ conditions for 3 days and analysed for GUS signals. Scale bar=500µm. **(e)** Split root assay showing pH_Ext_ mediated root growth inhibition as a systemic as well as local response. Bifurcated roots from seedlings grown at pH_5.8_ were transferred to different pH_Ext_ conditions (as indicated) and analysis was done was done after 3 days of treatment. (**f)** Relative PRL showing local and systemic RGI during acidic and alkaline pH_Ext_ conditions. PRL was normalised against optimum pH_5.8_ and average values of 10 independent replicates were plotted.

A very recent report suggested that the rhizospheric pH can accordingly influence apoplastic pH resulting into cellular expansion in roots (**Barbez et al., 2017**). To test whether root cell elongation is the major factor resulting into the observed RGI in response to rhizospheric pH, we analysed the epidermal cell length in roots subjected to different pH conditions. Our results validated the previous report where root epidermal cell length was negatively correlated (r=-0.9996) with the rhizospheric pH over the tested pH_Ext_ range (**Figure 1c, Figure S2**). However, the contradicting pH_3.8_ mediated RGI in spite of root cell elongation failed to justify the notion that cell elongation alone can result into the observed PRL over the tested pH_Ext_ range. Knowing that cell size and meristem activity both contribute to the root length, we tested the meristematic activity utilizing expression of cyclin B1-GUS (CYCB1;1:GUS) Arabidopsis reporter lines. Allowing the visualisation of cells entering the G2-M transition, GUS signals in CYCB1;1:GUS revealed a significant impact of rhizospheric pH on root apical meristem (RAM) activity (**Figure 1d**). The GUS signals were highest at pH_5.8_ with steady reduction towards both pH_3.8_ as well as pH_7.8_ well correlating with the observed PRL. These observations suggest a significant decrease in the meristem activity upon pH_Ext_ change from pH_5.8_. Therefore, the inhibited PRL during extreme pH_Ext_ conditions seems to be a concerted action of both cell size and cell division.

### Rhizospheric pH mediated RGI is a local as well as systemic response

In another set of experiments, we utilized split root assays where anchored roots from a single plant were allowed to grow on media in combinations of pH_5.8_ with pH_3.8_, pH_5.8_ and pH_7.8_. The root growth analysis with these experiments revealed a differential pattern of RGI. The split roots growing on pH_5.8_ (control conditions) show relative PRL ratios of 1. When these splits roots assays were performed in combination of pH_3.8_ and pH_5.8_, a better root growth was observed at pH_3.8_ along with inhibited root length at pH_5.8_. This resulted in the high relative PRL ratios (>1) when compared to the previous combination (**Figure 1e,f**). In contrast, when the combination of pH_5.8_ and pH_7.8_ was analysed, root growth inhibition was observed at pH_7.8_ without affecting the roots growth at pH_5.8_ resulting in lower PRL ratios (<1). These results suggested that while the acidic pH_3.8_ has systemic effect on root growth, the pH_7.8_ mediated RGI is purely a local response.

### Auxin gradient modulates RGI under fluctuating rhizospheric pH

Auxin is known to regulate apoplastic pH and therefore, the cell expansion in response to rhizospheric pH (**Barbez et al., 2017**). Further, the reversible nature of auxin mediated RGI (**Fendrych et al., 2018**) similar to as observed in our experiments (**Figure 1b**), led us to test the role of auxin in pH_Ext_ mediated RGI. In order to test this hypothesis, we first analysed how pH_Ext_ impacts the growth behaviour of roots towards gravity stimulus. For this we changed the gravity vector by 90-degree angle for the seedlings growing on different pH_Ext_ conditions and analysed the root angle formed with the initial gravity vector. Our observations revealed an increased gravitropic response with increasing pH_Ext_ (**Figure 2**) strengthening the hypothesis that auxin is certainly involved in plant’s response against pH_Ext_.

**Figure 2.**
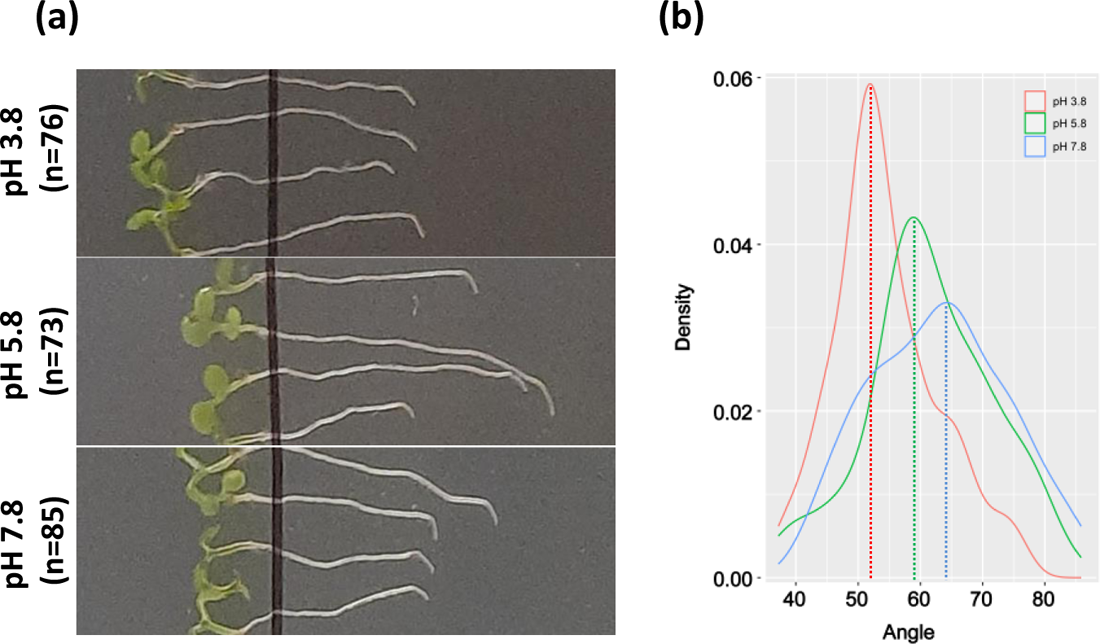
Rhizospheric pH modulates gravitropic response in Col-0. **(a)** Representative image showing differential gravitropic response in Col-0 against different pH_Ext_ conditions. **(b)** Density plot showing root angle formed in Col-0 against different pH_Ext_ conditions. Seedlings germinated on ½ MS media were subjected to 90 degree gravity vector change and root angle was recorded 10 hours after changing the growth orientation. Dotted line represents the maximum representation of root angle formed in response of gravity vector change w.r.t. different rhizospheric pH.

After knowing this possible role of auxin in pH_Ext_ mediated RGI, we tested if there is any change in auxin distribution at root tip subjected to different pH_Ext_ conditions. To record such dynamic fluctuations in root auxin, we subjected DR5rev::GFP (auxin responsive synthetic promoter driving GFP) seedlings to different pH_Ext_ conditions and analysed the fluorescence signals. Interestingly, we observed that DR5rev::GFP seedlings grown on pH_3.8_ displayed lowest fluorescence signal while this signal intensity was gradually increased on increasing pH_Ext_ with a strong correlation (r=0.93) (**Figure 3a**). Considering the possibility of any impact of pH_Ext_ on stability of GFP itself, we re-confirmed the auxin pattern with DII-VENUS, where, in contrast to DR5rev::GFP this reporter inversely correlates with the auxin accumulation and signalling (**Brunoud et al., 2012**). Now in this case, we observed highest fluorescence signals at pH_3.8_ with a gradual decrease with increasing pH_Ext_ conditions again following linear but a negative correlation (r=-0.96) (**Figure 3b**). These results with DR5rev::GFP and DII-VENUS confirmed that the auxin maxima at root tip is being regulated by pH_Ext_ with low auxin abundance at pH_3.8_ and high auxin at pH_7.8_. Knowing that auxin inhibits root growth both at its higher as well as lower concentrations (**Evans et al., 1994**), the RGI during different pH_Ext_ conditions can well be explained with the auxin abundance at root tip.

**Figure 3.**
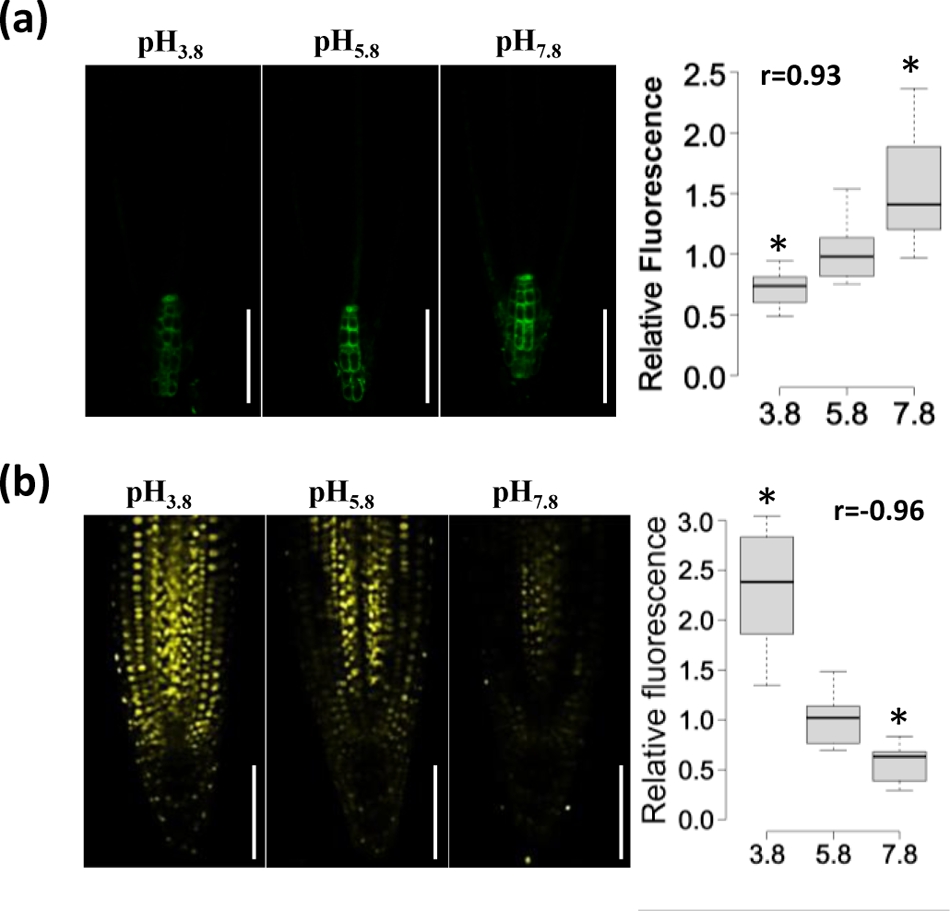
Auxin maxima at root tip explains RGI against pH_Ext_. Representative images and quantitative analysis of fluorescence in **(a)** DR5rev::GFP and **(b)** DII:VENUS seedlings subjected to different pH_Ext_ conditions. DR5rev::GFP or DII:VENUS seedlings were transferred to indicated pH_Ext_ for 3 days and then visualized under confocal microscope. n≥17 and n≥7 for DR5rev::GFP and DII:VENUS, respectively. Relative fluorescence was plotted after normalizing with average fluorescence values at pH_5.8_. Correlation coefficient (r) was calculated between average fluorescence and pH_Ext_ values, p<0.05, Scale bar=100µm.

### Auxin biosynthesis and catabolism is associated with pH_Ext_ mediated RGI

To check the direct involvement of auxin we observed RGI on different pH_Ext_ conditions supplemented with a gradient of NAA (a synthetic auxin). To our observations, Lower concentrations of NAA treatment rescued RGI on pH_3.8_ while seedlings on pH_7.8_ with NAA treatment showed hypersensitive RGI as compared to pH_5.8_ (**Figure 4a**). Considering these observations together, we proposed that low endogenous auxin at root tip on pH_3.8_ or higher endogenous auxin on pH_7.8_, inhibits RAM activity resulting into the observed RGI. In order to further explore the mode of auxin abundance in root tip, we utilized pharmacological inhibitors for the polar auxin transport and auxin biosynthesis. Arabidopsis seedlings subjected to either NPA or TIBA resulted into as similar pH_Ext_ mediated RGI pattern as to NAA treatment. Seedlings with NPA and TIBA treatment showed a hyposensitive RGI at pH_3.8_ but exaggerated RGI on pH_7.8_ conditions (**Figure 4b-c**). As NPA and TIBA both are auxin efflux inhibitors (**Muday et al., 2002; Abas et al., 2021**), it seems to be cellular auxin scarcity/toxicity at RAM during severe pH_Ext_ conditions resulting into the observed RGI. Therefore, the localized auxin maxima at root tip seem to be the modulator of RGI during severe pH_Ext_ conditions. To further validate these results, we analysed the abundance of PIN transporters (using in roots subjected to different pH conditions. Comparative studies with pPIN1::PIN1:GFP and pPIN2::PIN2:GFP revealed decreasing abundance of PIN1 and PIN2 towards either side of the rhizospheric pH from pH_5.8_ (**Figure 4d-e**). Interestingly, PIN7 (pPIN7::PIN7:GFP) showed a linear pattern of abundance throughout the pH_Ext_ range (**Figure 4f**). PIN7 abundance in stele region showed a linear decrease towards increasing value of pH_Ext_ while PIN7 abundance in columella showed linear increase towards increasing pH_Ext_ value. As PIN proteins facilitate auxin export out of the cell, increasing abundance of PIN7 in columella region with increasing pH_Ext_ value seems to be an adaptive response for distributing the over-accumulated auxin to reduce its detrimental effect on root meristem. Further, the decreasing abundance of PIN7 in stele region towards increasing pH_Ext_ again help in reducing further auxin accumulation in root meristem through inhibiting the acropetal auxin transport.

**Figure 4.**
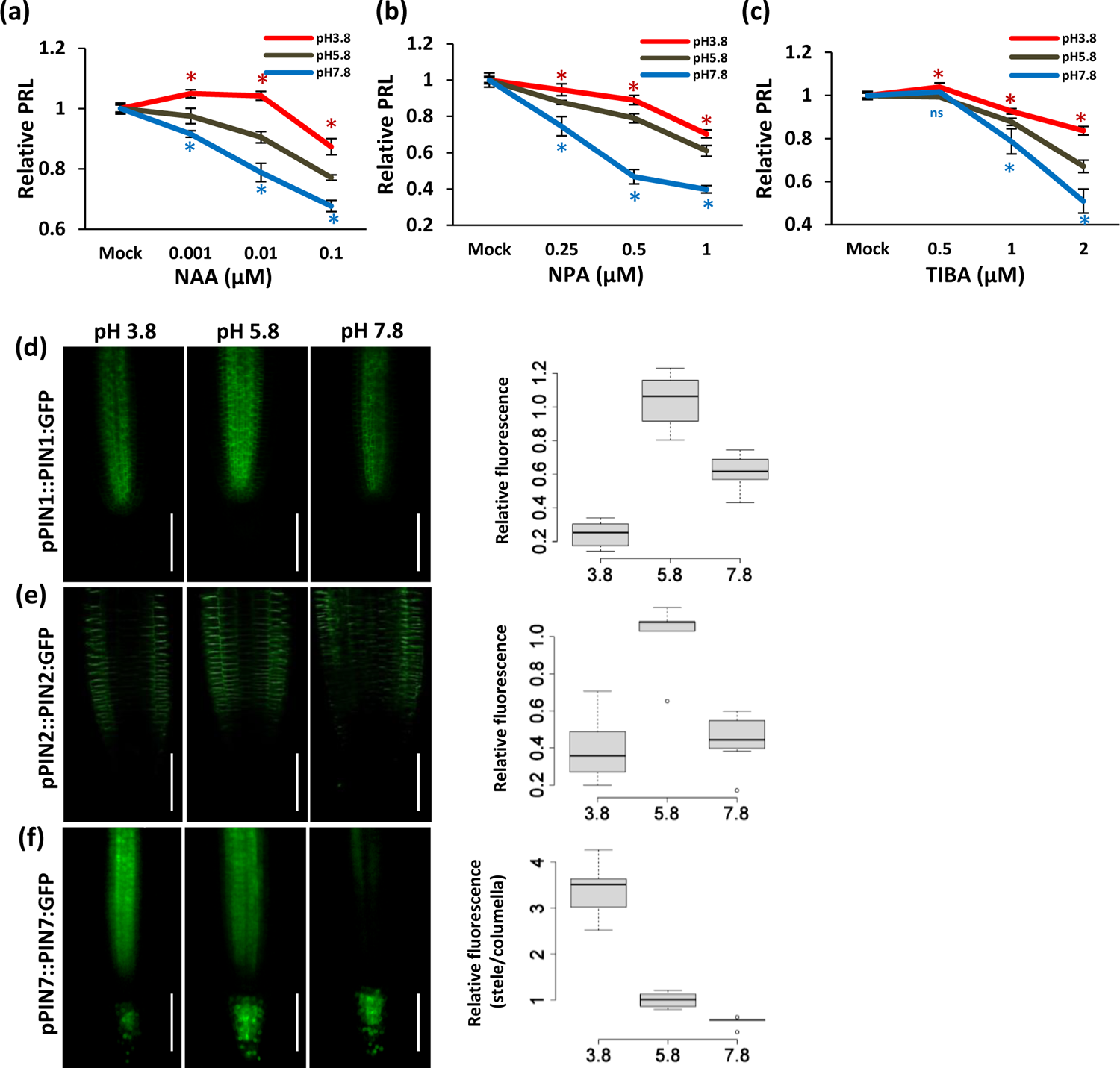
Localised auxin maxima at root tip is associated with pH_Ext_ mediated RGI. Relative root growth analysis (Col-0) on different pH_Ext_ conditions supplemented with indicated concentrations of **(a)** NAA (n≥20), **(b)** NPA (n≥20), **(c)** TIBA (n≥20). PRL was quantified after 5 days of treatment and normalized with mock (i.e. 0µM NAA/NPA/TIBA) for plotting. PRL was quantified after 5 days of treatment and normalized with PRL at pH_5.8_ for plotting. Student’s *t-*test was applied for significance analysis, p<0.05. Representative images and fluorescence quantification of **(d)** pPIN1:PIN1:GFP, **(e)** pPIN2::PIN2:GFP and **(f)** pPIN7::PIN7:GFP subjected to different pH_Ext_ conditions. Uniformly germinated seedlings were transferred to different pH conditions and fluoresce signals were analysed after 3 days of treatment. Relative fluorescence was plotted after normalizing with average fluorescence values at pH_5.8_. Correlation coefficient (r) was calculated between average fluorescence and pH_Ext_ values, p<0.05, Scale bar=100µm, n≥5.

Interestingly, seedlings treated with either PPBo or BBo (inhibitors of YUCCA mediated auxin biosynthesis) individually show an inverse pattern as compared to NAA/NPA/TIBA treatments. Seedlings growing on pH_3.8_ showed hypersensitive RGI while seedlings growing on pH_7.8_ were hypo-sensitive to pH_Ext_ mediated RGI when supplemented with BBo or PPBo (**Figure 5a-b**). This suggested that although auxin abundance is strictly associated with pH_Ext_ mediated RGI, YUCCA mediated auxin biosynthesis may not have a specific role to play. We then went on to check if TAA1/TAR pathway is associated with pH_Ext_ mediated auxin accumulation. For this, we analysed RGI in seedlings growing on different pH_Ext_ supplemented with L-kynurenine [kyn, an inhibitor of TAA1/TAR pathway, (**He et al., 2011)**]. Interestingly, seedlings treated with kyn were found to be equally sensitive to pH_3.8_ mediated RGI as to pH_5.8_. However, kyn treatment rescued pH_7.8_ mediated RGI in a dose dependent manner though at lower concentrations (**Figure 5c)**. Therefore, TAA1/TAR pathway seems to be associated with pH_Ext_ mediated auxin abundance by regulating its biosynthesis towards alkaline pH_Ext_ conditions. Furthermore, role of auxin receptor gene transport inhibitor response 1 (TIR1, a F-box protein) was checked to ascertain if auxin perception/signalling is required for pH mediated RGI. To our observations, *tir1-1* mutant responded as similar as Col-0 against pH_3.8_ as well as pH_7.8_ conditions, suggesting the RGI regulation is TIR1 independent **(Figure S3)**. Overall, we confirm that auxin play an indispensable role in pH_Ext_ mediated RGI though in a TIR1 independent manner.

**Figure 5.**
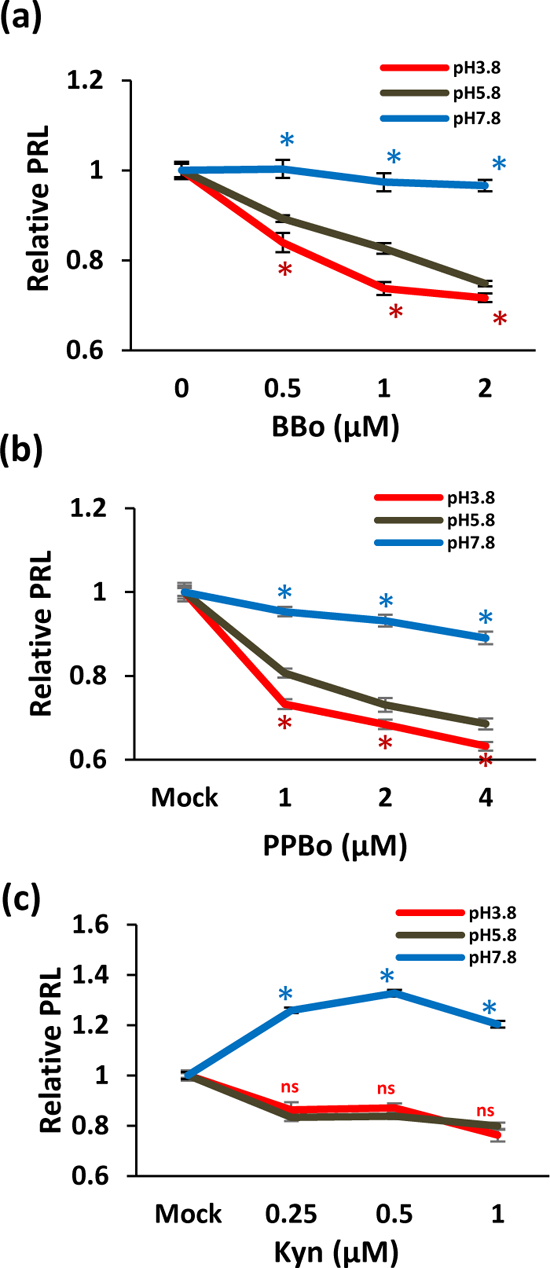
Effect of auxin biosynthesis inhibitors on pH_Ext_ mediated RGI. Relative root growth analysis (Col-0) on different pH_Ext_ conditions supplemented with indicated concentrations of (a) BBo (n≥20), **(b)** PPBo (n≥25) and **(c)** kyn (n≥20). PRL was quantified after 5 days of treatment and normalized with mock (i.e. 0µM BBo) for plotting. Asterisks marks the significant difference with student’s *t-*test, p<0.05.

### Jasmonates regulate auxin catabolism in a pH_Ext_ dependent manner

Since, auxin maxima can be regulated via, auxin biosynthesis, transport or catabolism. Our results excluded the involvement of auxin transport in pH_Ext_ mediated RGI, however, auxin biosynthesis through TAA1/TAR pathway seems to be involved. Next, we studied the genes involved in the auxin catabolism under varying pH conditions. Previous report on the microarray data for acidic pH conditions suggested up-regulation of GRETCHEN HAGEN 3 (GH3) family genes such as *GH3.1* and *GH3.3* (**Lager et al., 2010**). These enzymes are known to catabolize auxin by conjugating it to different amino acids (**Hayashi et al., 2021**). We studied the expression of all GH3 family members on different pH_Ext_ conditions. In agreement with the previous report, our analysis confirmed upregulation of both *GH3.1* and *GH3.3* at pH_3.8_ w.r.t. pH_5.8_ (**Figure 6a**). Interestingly, expression of *GH3.1* remained unchanged at pH_7.8_ while *GH3.3* was found induced at both, pH_3.8_ as well as pH_7.8_. Notably, we observed inverse correlation (r=-0.98) between expression of *GH3.9, GH3.10, GH3.11* and *GH3.17* with the pH_Ext_ value.

**Figure 6.**
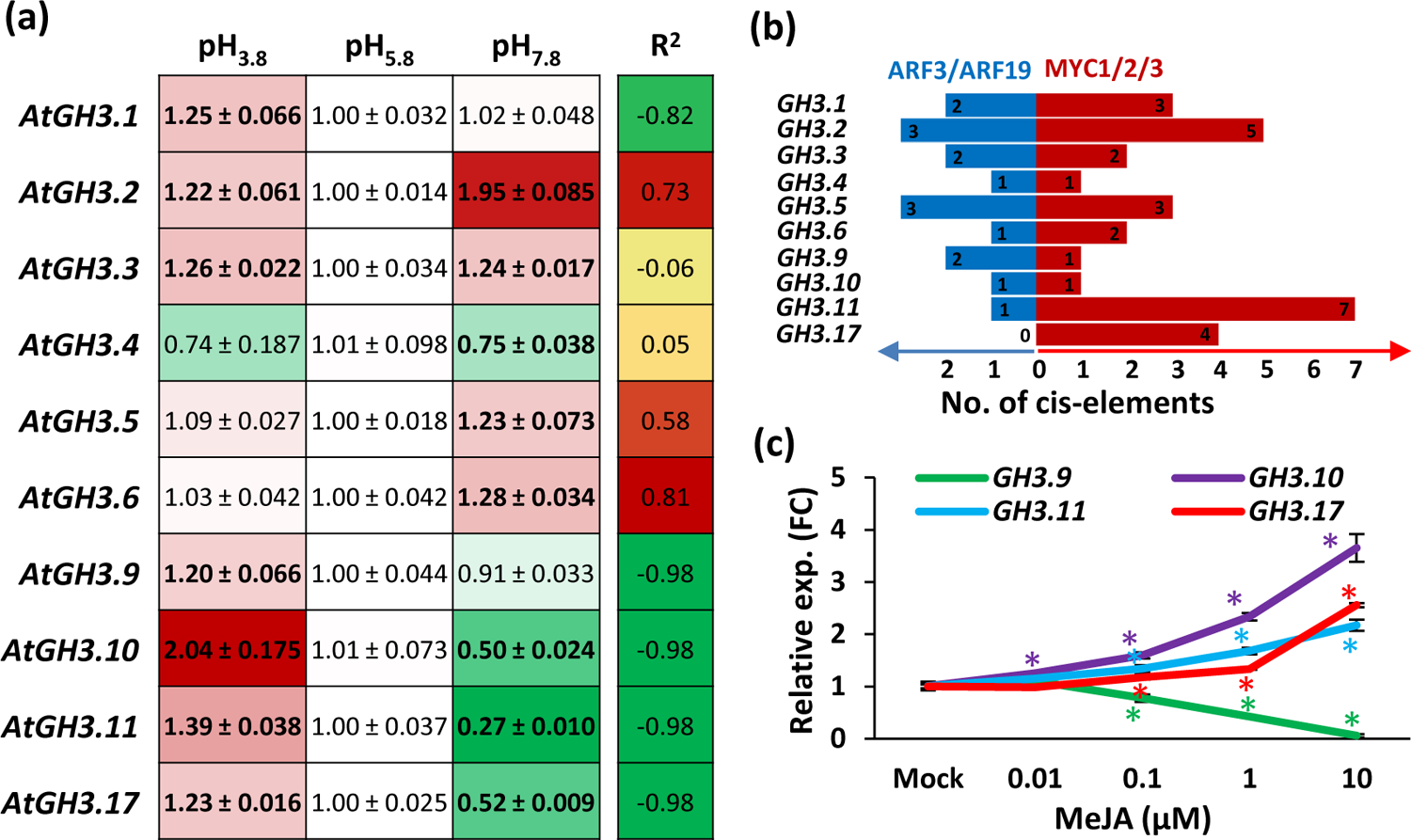
Jasmonate dependent auxin catabolism is associated with pH_Ext_ mediated RGI. **(a)** Heat-map showing transcriptional behaviour of *GH3* family genes (relative expression levels, FC) during different pH_Ext_ conditions in roots. Expression was normalized with expression at pH_5.8_ after 8 days of the treatment. Bold entries indicate the significant change as compared to pH_5.8_. n≥3, students *t-*test, p≤0.05. **(b)** Number of ARF3/ARF19 and MYC1/2/3 TFs binding sites in the promoter region of each *GH3* family genes to show the cis-element enrichment. Promoter was identified as 2kb upstream region of the transcription start site using PlantPan2.0. (http://plantpan2.itps.ncku.edu.tw/). Numbers in the horizontal bars indicate number of respective cis-elements. **(b)** Line graph showing transcriptional behaviour of *GH3.9*, *GH3.10*, *GH3.11* and *GH3.17* on MeJA treatment. Each datapoint represent average fold change values of n≥3 with SE among the replicates. Asterisks indicate the significant change w.r.t. the expression at mock treatment.

Considering the previously published reports, GH3.10 and GH3.11 (JAR1) do not catabolize auxin but are known to activate Jasmonic acid (JA) by converting it to JA-Ile (JA-Isoleucine) (**Staswick and Tiryaki, 2004; Staswick et al., 2005; Chen et al., 2017; Delfin et al., 2022**). The promoter analysis of GH3 family genes suggests enrichment of JA associated cis-elements regulated by transcription factors like MYC1/2/3 (**Figure 6b**). Further, exogenous application of MeJA (methyl jasmonate) on the Arabidopsis seedlings, show dose dependent induction of *GH3.10*, *GH3.11* and *GH3.17* though *GH3.9* was downregulated **(Figure 6c)**. This data suggested a strong negative association of JA with auxin abundance. In order to further comment on involvement of JA in auxin regulated RGI against varying pH_Ext_ conditions, we analysed the influence of MeJA treatment on pH_Ext_ mediated RGI. Our results supported almost linear nature of MeJA mediated RGI at pH_5.8._ In contrast, seedlings were relatively insensitive to the treatment at pH_3.8_ and pH_7.8_ (**Figure 7a**). As Jas9:VENUS reporter lines can be used as biosensor for active JA, we utilized this reporter to probe spatial changes in JA abundance on varying pH_Ext_ conditions **(Larrieu et al., 2015)**. We observed lowest Jas9:VENUS fluorescence signals on pH_3.8_ with increasing fluorescence towards pH_7.8_ following a strong linear correlation (r=0.97) with the increased pH_Ext_ (**Figure 7b-c**). As Jas9:VENUS fluorescence follows an inverse pattern with JA abundance, these results revealed high JA accumulation on pH_3.8_ and lowest accumulation on pH_7.8_ with a linear change along the entire pH_Ext_ range. Further, GH3.17 is known to catabolise auxin, MeJA mediated induction of *GH3.17* support the notion that JAs work upstream of auxin and modulate auxin abundance which in turn regulate RGI in response to pH_Ext_.

**Figure 7.**
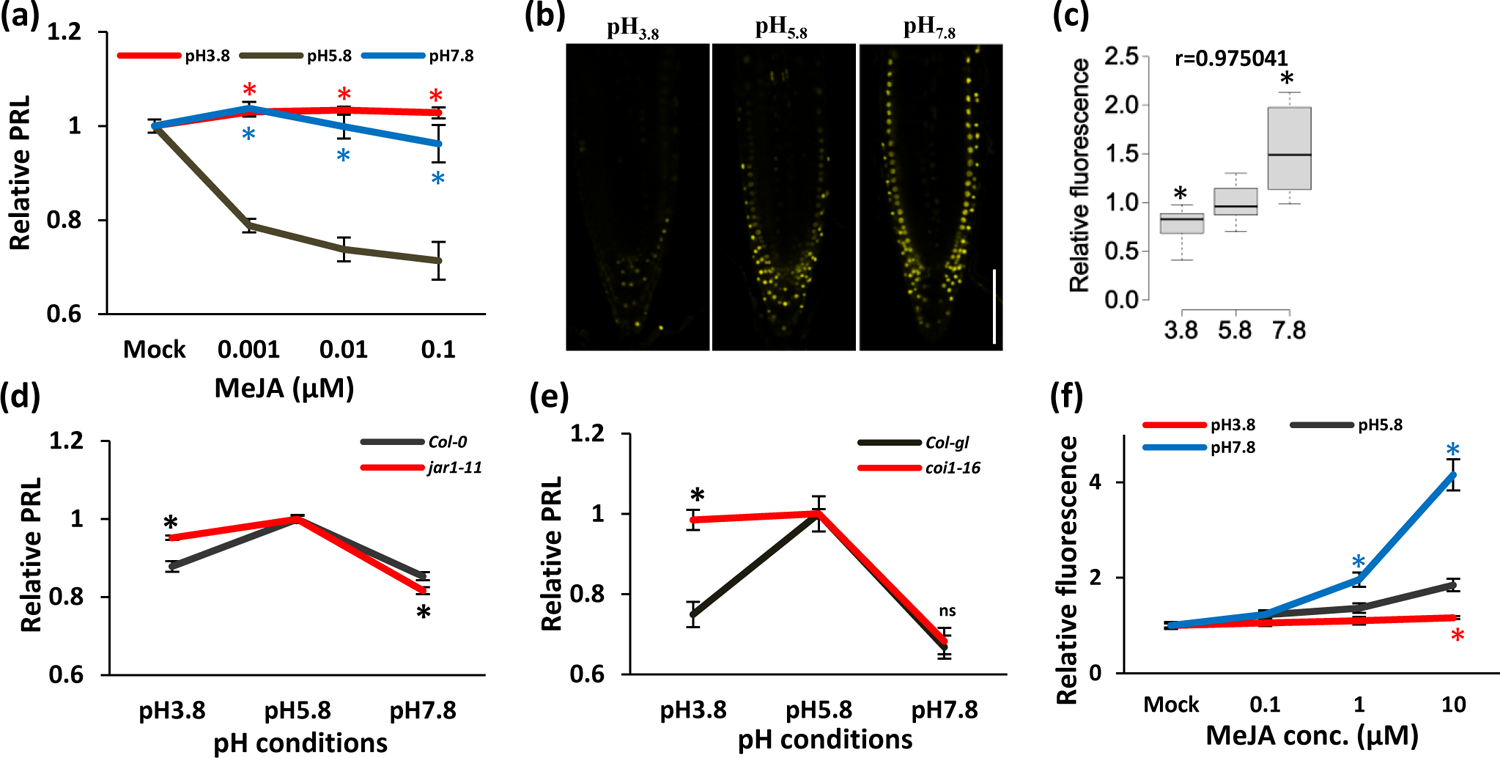
Jasmonate signaling inversely explains auxin mediated RGI against pH_Ext_. **(a)** Relative PRL of Col-0 at different pH conditions supplemented with a gradient of MeJA. PRL was normalized to mock treatment at the respective pH conditions and average values were plotted with SE among the replicates. Significance was analysed w.r.t. relative PRL at pH_5.8_ with the respective MeJA concentration. **(b)** Representative images showing effect of pH_Ext_ on Jas9:VENUS. **(c)** Quantitative analysis of Jas9:VENUS florescence subjected to different pH_Ext_ conditions. Fluorescence was normalized with pH_5.8_, x-axis represents the pH_Ext_ value, n≥14, Asterisks indicate the significant difference as compared to pH_5.8_. Correlation coefficient (r) was calculated between average fluorescence and pH_Ext_ values. Relative PRL analysis of **(d)** Col-0 and *jar1-11* (n≥25) and **(e)** col-gl and *coi1-16* (n≥12) under indicated pH_Ext_ conditions. PRL was quantified after 5 days of treatment and normalized with PRL at pH_5.8_ for plotting each graph. Each datapoint indicate the average relative PRL values with SE among the replicates. **(f)** Relative fluorescence values of DII:VENUS under indicated pH_Ext_ conditions supplemented with different concentrations of MeJA. PRL and fluorescence levels were analysed after 5 days of treatment and normalized with Mock (i.e. 0µM MeJA) for plotting for each pH_Ext_ condition individually. Asterisks marks the significant different with student’s t-test, p<0.05, n≥5. Asterisks marks the significant different with student’s *t-*test, p<0.05. Scale bar=100µm.

### JA antagonise auxin response during pH_Ext_ mediated root growth inhibition

*JAR1* (*GH3.11*) expression was found strongly linked to the pH_Ext_ justifying JA abundance during the entire range of pH_Ext_. Therefore, we tested the involvement of JAs in pH mediated RGI using *jar1-11* mutant which is a JA-Ile deficient mutant with a reduced sensitivity to JA mediated root growth inhibition (**Suza and Staswick, 2008**). Our observations marked *jar1-11* relatively insensitive to pH_3.8_ and hypersensitive to pH_7.8_ mediated RGI (**Figure 7d**). These observations confirmed the dependency of pH_Ext_ mediated RGI on JAs over the entire pH_Ext_ range. Now, to answer whether JA perception is required for pH_Ext_ mediated RGI, we analysed growth behaviour of *coi1-16* (a JA perception mutant), on different pH_Ext_ conditions (**Ellis and Turner, 2002**). We observed that while *coi1-16* behaved similar to Col-gl (WT) at pH_7.8_ in terms of RGI, it was specifically found insensitive at pH_3.8_ mediated RGI (**Figure 7e**).

Identification of MYC1/2/3 binding sites in promoters of *GH3* gene family members (**Figure 6b**) along with an inverse correlation of auxin and JA abundance during entire range of pH_Ext_ conditions, suggested a potential cross-talk between these two phytohormones. To elucidate this cross-talk, we subjected DII:VENUS lines to different pH_Ext_ supplemented with a gradient of MeJA. Our analysis on DII:VENUS show linear increase in fluorescence with increased MeJA concentration on pH_5.8_. Whereas, least change in fluorescence on pH_3.8_ and hyper-accumulation of DII:VENUS fluorescence was observed on pH_7.8_ condition in a dose dependent manner (**Figure 7f**). These observations suggested an antagonistic effect of MeJA treatment on auxin accumulation. At pH_3.8_ high JA content is suggested to lower down the auxin accumulation at root tip while at pH_7.8_ very low JA failed to inhibit auxin accumulation resulting into auxin toxicity. These observations further justified our previous results where a relatively insensitive phenotype was observed w.r.t. MeJA mediated RGI on pH_7.8_ (**Figure 7a**). While these results validated the pH_Ext_ mediated RGI as a function of JA regulated auxin accumulation, also extended impact of JAs on alkaline pH_Ext_ mediated RGI suggesting that these two hormones regulate RGI over the entire range of pH_Ext_.

## Discussion

Considering the importance of rhizospheric pH in determining the plants fitness and yield, elucidation of the mechanism of pH_Ext_ perception and its response finds a great importance. Very recently, RGI was marked as characteristic feature of plants response against extreme rhizospheric pH while the factors associated with such RGI are the topics of active research (**Tsai and Schmidt, 2021; Bailey et al., 2022**). Further, it was also delineated that acidic rhizospheric pH positively impacts on root cell elongation but in spite of cellular elongation root growth was inhibited which failed to support the idea of involvement of acid growth theory alone in pH_Ext_ mediated RGI (**Bailey et al., 2022; Barbez et al., 2017)**. Loss of meristematic activity on extreme pH conditions while justifying the root growth on entire pH range, also suggested a complex molecular response rather just being a physical phenomenon like of acid growth theory. The differential mode of RGI (local as well as systemic as on pH_7.8_ and pH_3.8_, respectively) further added to its complexity and suggested involvement of multiple factors. One of these factors was easily identifiable by the reversible nature of RGI which aligns well with the auxin response (**Fendrych et al., 2018**). Again, transcriptomic induction of auxin related genes strengthened involvement of auxin in the response (**Bailey et al., 2022**). The linear increase in auxin abundance/signalling (as indicated by DR5rev::GFP and DII:VENUS) on increasing pH_Ext_ value forms direct evidences for involvement of auxin in pH_Ext_ mediated RGI. Further, the known facts for root growth inhibition by virtue of both low and high auxin abundance strengthened our results.

The analysis further extended our knowledge over the mode of auxin metabolism in response to rhizospheric pH as auxin transport inhibitors (NPA/TIBA) failed to rescue the RGI effectively on either side of the optimal pH_Ext_ rather they intensified the RGI on pH_7.8_ which was a similar phenotype as observed on auxin treatment itself (NAA, **Figure 4**). Similar phenotypic observations on PAT (polar auxin transport) inhibition were marked previously (**Duan et al., 2023**). As TIBA and NPA are auxin export inhibitors, our experiments suggested a localised toxicity of auxin during alkaline pH as the cause of RGI which is in accordance to the previous study. Our conclusion was further supported by the reduced abundance of PIN1, PIN2 and PIN7 (in stele) during strongly alkaline pH_Ext_ forms an adaptive response rather contributing to the auxin toxicity at root tip. In addition to this, increased abundance of PIN7 in columella towards increasing pH_Ext_ value again confirmed the adaptive nature of PIN proteins to reduce the auxin abundance at root meristem (**Figure 4**). Further, NAA treatment mediated rescue of RGI on pH_3.8_ confirmed the cause as low auxin abundance (**Figure 4**). While its hypersensitivity on pH_7.8_ due to auxin overaccumulation was confirmed with the previous results where auxin signalling deficient mutant *arf7arf19* showed better root growth on more alkaline pH_Ext_ as compared to pH_5.8_ (**Duan et al., 2023**).

Interestingly, TAA1/TAR and YUCCA both participate in the same branch of auxin biosynthesis (**Zhao, 2014**) but only TAA1/TAR were found to be influenced by pH_Ext_ suggesting that pH_Ext_ regulate auxin biosynthesis by inhibiting the rate limiting step of the pathway leaving role of YUCCA family inefficient. Rhizospheric pH_3.8_ seems to inhibit the TAA1/TAR pathway very efficiently leaving no room for further inhibition by inhibitors like kyn (**Figure 5**). The expression pattern of *GH3.17* participating in auxin catabolism (**Staswick et al., 2005**), strongly correlated (negatively) with the pH_Ext_ values and therefore, seems to play critical role in pH_Ext_ mediated auxin accumulation. Our analysis suggested that a concerted action of both TAA1/TAR mediated biosynthesis and GH3.17 mediated catabolism of auxin contributes to the final auxin levels in the roots as determined by the pH_Ext_ condition. Further, having no impact of *TIR1* mutation on RGI suggested an TIR1 independent pathway regulating pH_Ext_ meditated RGI (**Figure S3**). The notion of TIR1 independent pathway regulating pH_Ext_ mediated RGI was further supported by the reversible nature of the RGI as observed previously that auxin may regulate RGI in a reversible manner (**Fendrych et al., 2018**) and can be TIR1 independent (**Monshausen et al., 2011**).

Further, the qRT-PCR based expression analysis of *GH3.10, GH3.11* and *GH3.17* on different pH_Ext_ conditions and MeJA treatment in addition of confocal microscopy based validation of induced DII:VENUS signals on MeJA treatment marked the crosstalk between JA and auxin signaling (**Figure 6**, **Figure 7f**). As auxin mediated responses are largely local while JAs are known for systemic mode of response, we proposed that these two factors are responsible for the pH_Ext_ mediated RGI. Our results, with increasing concentration of JA towards low pH (**Figure 7b-c**) justified the systemic response where split roots experiencing pH_3.8_ may exert an inhibitory effect on roots growing on pH_5.8_, when attached to the same seedling. Further, pH_3.8_ also resulted into insensitive behaviour of seedlings towards MeJA mediated RGI (**Figure 7a**). The better growth of root experiencing pH_3.8_ (both as in **Figure 1e** and **Figure 7a**) in spite of having higher JA can be explained by the adaptive nature of JAs during stress conditions as proved earlier with our study where higher intrinsic JA can result into JA insensitivity in plants (**Singh et al., 2020**). On the other hand, decreasing JA abundance towards pH_7.8_ resulted into vanished JAs mediated systemic response while the increasing auxin accumulation leads to the root growth inhibition as a local response. Though pH_Ext_ sensing by plants need further investigation, we proved that these two phytohormones seems to play the central role in pH_Ext_ mediated RGI.

With the current understanding we propose that certain trans-acting factors [as speculated earlier (**Tsai and Schmidt, 2021**)] regulate GH3.10 and GH3.11 (JAR1) resulting into the appropriate change in JA abundance in response to rhizospheric pH fluctuations (**Figure 8**). Subsequently, JAs modulate expression of *GH3.17* gene which finally dictate the auxin maxima supporting the root length. Therefore, our study delineated the novel crosstalk between jasmonates and auxin in the regulation of pH_Ext_ mediated RGI further extending our understanding for soil pH perception and downstream response by plants.

**Figure 8.**
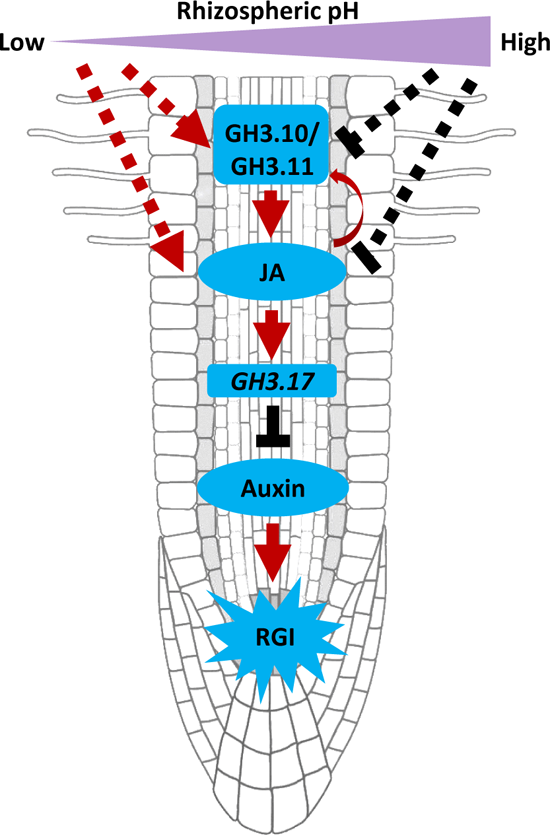
A working model explaining regulation of RGI during differential rhizospheric pH. Rhizospheric pH regulate JA accumulation where acidic and alkaline pH_Ext_ supports high and low JA content, respectively, possibly in a GH3.10 and GH3.11 (JAR1) dependent manner. On the other hand JA and JAR1 forms a positive feedback loop regulating each other. This accumulated JA then positively regulate auxin catabolism related gene transcripts (*GH3.17*) which in turn negatively dictate auxin maxima in roots. Finally, this differential auxin maxima in roots results into pH_Ext_ mediated RGI based on auxin deficiency or toxicity during acidic or alkaline pH_Ext_ conditions, respectively.

## Acknowledgements

We thank Jitender Giri and Ashverya Laxmi (National Institute of Plant Genome Research, New Delhi, India); and Saikat Bhattacharjee (Regional Centre for Biotechnology, Faridabad, India) for sharing materials. We also thank Confocal facility, NABI, India for imaging. Part of this research is funded by the MK Bhan Young Researcher Fellowship, DBT, India to APS and NABI CORE grant to AKP.

## Authors’ contributions

A.P.S. and A.K.P. conceived the project, designed experiments, analysed data and wrote the manuscript. A.P.S. and R.K. performed all the experiments.

## Supporting Information

**Supplementary Figure S1**. Phenotypic behaviour of Col-0 over the pH_Ext_ range.

**Supplementary Figure S2**. Impact of pH_Ext_ on root cell length.

**Supplementary Figure S3**. Effect of pHExt on root growth inhibition in Col-0 and *tir1-1* mutant.

**Supplementary Table S1**. List of primers used in the study.

